# Age-related cerebellar genetic, neuronal and functional impairments are reversed by specific magnetic stimulation protocols

**DOI:** 10.64898/2026.03.24.713842

**Authors:** Aurélien Fauquier, Tom Dufor, Natalie Morellini, Mohamed Doulazmi, Jean Mariani, Ann M Lohof, Rachel M Sherrard

**Affiliations:** Sorbonne Université, CNRS UMR8263 and INSERM U1345, Development, Adaptation and Ageing, Paris, France

**Keywords:** LI-rTMS, ageing, cerebellum, Purkinje cells, spatial memory, dendrites, magnetic fields

## Abstract

Age-related cognitive decline reflects progressive atrophic changes that advance through broad neural networks. There is no effective treatment. However, brain ageing is *not homogenous,* so treating the earliest-affected circuits may be successful in reversing and/or preventing ongoing neuronal atrophy and therefore cognitive decline. Repetitive transcranial magnetic stimulation (rTMS), a non-invasive technique that modulates cortical excitability, induces activity-dependent neuronal plasticity. Here we investigate short- and long-term effects of low intensity rTMS (LI-rTMS) on the cerebellum, which is adversely affected early during ageing. With age, cerebellar genes related to inflammation are strongly upregulated, whereas processes of synaptic-maintenance are reduced. Both abnormalities are rapidly corrected by LI-rTMS in a protocol-dependent manner. In parallel, LI-rTMS increases neuronal spine density and dendritic complexity, in association with improved spatial memory in both young adult *and* aged mice. These responses of the ageing cerebellum to low-intensity magnetic stimulation are extremely encouraging for treating age-related cognitive decline, but reinforce that appropriate stimulation parameters must be identified for effective treatment.

## INTRODUCTION

Age-related cognitive decline imposes profound social and economic burdens, even before the onset of dementia with its associated loss of autonomy for elderly people ^1^. This decline is driven by complex cellular changes that cause synaptic dysfunction and progressive disruption extending through large-scale neural networks and reducing complex cognitive function ^2,3^. Currently there are no treatments that can reverse age-related neurodegeneration; at best a healthy life-style or psychomotor stimulation show potential for preserving physical and cognitive reserve ^4^. However, to repair dysfunctional brain circuits more concentrated activity-dependent neural plasticity, such as that induced by non-invasive brain stimulation (NIBS), is required. One NIBS method, transcranial magnetic stimulation (TMS), is a promising alternative for activating neural plasticity and mitigating age-related cognitive decline. Repetitive TMS (rTMS) alters cortical excitability, inducing neuroplastic changes, with established efficacy in the treatment of several neurological and psychiatric disorders ^5,6^ although effects are diminished in the aged brain ^7,8^. However, clinical outcomes are variable as the effects of magnetic stimulation depend on the structure and network connectivity of the brain region being stimulated^9^ at both functional and gene expression levels ^10,11^. Since age-related neuronal changes do not occur homogenously within the brain ^12^, treatments supporting function of early-affected circuits may prevent the expansion of dysfunction to other brain areas. Thus, outcomes of rTMS application to different brain regions need to be better understood, in order to optimise treatments for age-related cognitive decline.

The cerebellum is among the brain regions affected early during normal aging, and is attracting increasing attention for intervention because of its involvement in specific cognitive and emotive processes, in addition to long-established roles in motor coordination and balance ^13^. It is involved in age-related motor and cognitive decline ^14,15^ through its extensive cerebello-cerebral connectivity ^16^. Cerebellar information processing is greatly compromised with advancing age ^17^ through atrophy ^18^, altered dendritic arborization of its central macro-neuron, the Purkinje cell ^19^ and progressive neuronal loss ^17,20^. White matter atrophy also occurs, particularly in the superior cerebellar peduncle (cerebello-efferent axons), in parallel with decreasing function ^14,21,22^. Although cerebellar rTMS modulates its cerebro-cerebellar neuronal circuits to alter motor and cognitive function ^23,24,25^, the effects of magnetic stimulation on the cerebellum *per se*, which underlie these network changes, remain poorly defined. Moreover, outcomes from cerebellar rTMS cannot be predicted from studies on cortical stimulation, given the completely different structures of cerebellar and cerebral cortices and the particular vulnerability of the cerebellum during ageing.

Clinically, rTMS is delivered at high-intensity (0.5-2T), but its application is limited by the need for high-tension power sources, safety concerns and restricted stimulation parameters. In contrast, low-intensity rTMS (LI-rTMS; <20mT) enables a broader range of frequencies and stimulation patterns, enhances neuronal plasticity ^26,27,28^, and modulates brain activity with excellent tolerance ^29^. Its therapeutic potential is clear ^30,31,32^, due to described cellular mechanisms of reduced neuroinflammation ^33^, increased intracellular calcium and brain-derived neurotrophic factor (BDNF), altered gene expression, and the removal of aberrant synaptic connections in cortical and subcortical circuits ^26,27,34,35,36^. This low-intensity stimulation approach also runs on normal domestic electricity, opening the potential for home treatment. However, its specific impact on brain aging remains to be described.

In the present study, we evaluated the effects of LI-rTMS on the ageing brain, specifically targeting the cerebellum. We applied LI-rTMS to the cerebellum of young adult and aged mice, assessing the effect on gene expression profiles using RNA-Seq and qPCR as well as cerebellar structure and function to assess whether any genetic changes were associated with reversed age-related structural and behavioural deficits. Our findings show that ageing is associated with a marked upregulation of inflammation-related biological processes and a reduction in the synthesis of synaptic proteins in the cerebellum. These ageing related changes are normalised by some LI-rTMS protocols, in association with increased Purkinje cell dendritic and improved cognitive function.

## RESULTS

In this study we evaluated the effects of LI-rTMS on the mouse cerebellum of adult (3-5 months) and aged (17 months) mice. We observed that LI-rTMS rapidly alters cerebellar gene expression, so that abnormal expression linked to age was corrected, and this was followed by increased complexity of Purkinje cell dendritic tree morphology in association with improved spatial memory, a cerebellar-mediated behaviour.

### LI-rTMS Reverses Age-related changes in the Cerebellar Transcriptome

First, to obtain a baseline of cerebellar changes linked to ageing, we measured gene expression profiles by whole-tissue RNA sequencing in adult (4-months) and aged (17-months) cerebella from unstimulated (sham) mice. Ageing significantly altered the expression of 346 genes, with 76 downregulated and 270 upregulated (Fig. 1A). Gene Ontology (GO) analysis of these differentially expressed genes (DEGs) identified affected biological processes. GO analysis showed that ageing was associated with a marked increase in processes related to inflammation and immune system activity (Fig. 1B), for example immune system processes (p=3.46×10^-52^) and interleukin-6 production (p=1.25×10^-9^) and their many related sub-processes (Fig. 1C). GO analysis of the smaller group of downregulated DEGs, identified processes involving neuronal plasticity (Fig. 2A), e.g. synapse organization (p=8.76×10^-6^), neurogenesis (p=1.41×10^-5^), and synaptic membrane adhesion (p=8.8×10^-4^). This downregulation of genes regulating neuronal activity was then confirmed by qPCR. Gene expression associated with synaptic signalling and plasticity was downregulated in the aged vs adult cerebellum: specifically proteins involved in voltage gated potassium channels (*Kcnc3,* p = 0.002); postsynaptic scaffolding (*Psd95,* p = 0.0005; *Gphn,* p = 0.0005); neurotrophin signalling (*Ntrk2,* p = 0.0038); glutamate receptors (*Gria1,* p = 0.0004; *Syngap1,* p = 0.0244); and calcium regulation (*Calb1,* p = 0.0088; *Calb2,* p = 0.012; Fig. 2B). These changes indicate an age-related decline in synaptic integrity and plastic remodelling.

**Fig. 1.**
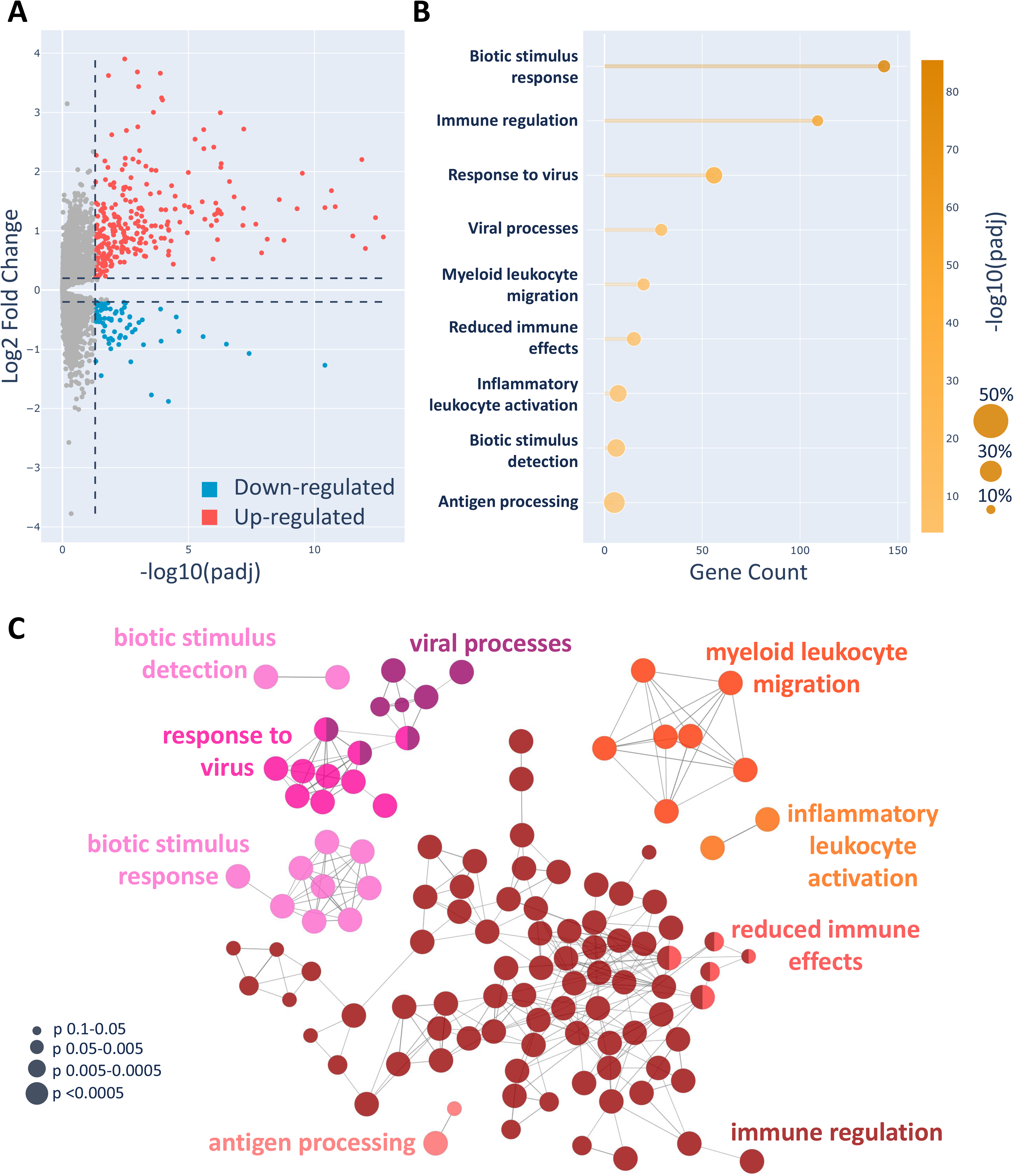
Ageing Increases Inflammation in the Cerebellum. **(A)** Volcano plot of differentially expressed genes in aged versus young-adult cerebellum. The vertical line marks the significance threshold (adjusted p value = pAdj < 0.05). Genes upregulated in ageing with a fold-change more than 0.2 (Log2FC > 0.2) are indicated in red, while downregulated genes (Log2FC < -0.2) are in blue. Grey dots represent genes that are not significantly changed or have minimal expression differences. **(B)** Gene Ontology (GO) analysis of biological processes increased by aging. The left vertical axis lists the top biological processes identified through Cytoscape analysis. Each pathway is associated with a gene count on the x-axis (number of altered genes in that pathway) and an adjusted p-value gradient, indicating the significance of enrichment. Bubble size reflects the proportion of genes in each biological process that are upregulated. **(C)** Gene Ontology (GO) map of biological processes upregulated by ageing. CLueGO results display a functionally-grouped network of pathways. The most significant pathway in each cluster is designated as the leading term. Node size represents pathway significance, and connecting lines indicate interactions between pathways.

**Fig. 2.**
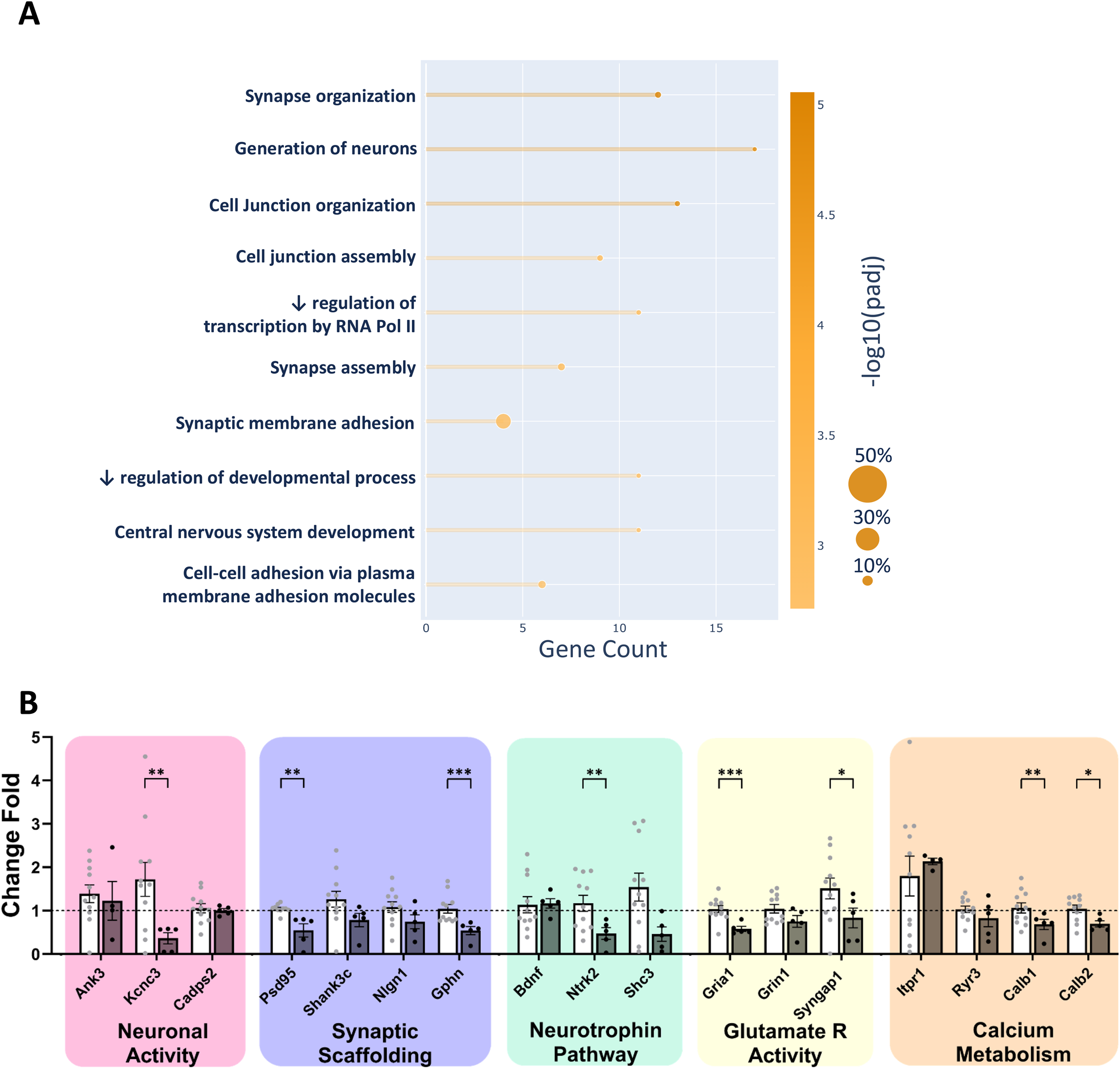
Aging Decreases Neuroplasticity in the Cerebellum. **(A)** GO analysis of biological processes decreased by ageing. The left vertical axis lists the top biological processes identified through GO. Each pathway is associated with a gene count on the x-axis (number of altered genes in that pathway) and an adjusted p-value gradient, indicating the significance of the gene set change. Bubble size reflects the proportion of genes in each biological process that are downregulated. **(B)** qPCR of some neuroplasticity genes that are important for cerebellar function. Data are expressed as fold change normalized to the adult sham group, according to the Pfaffl equation. The horizontal dotted line represents the mean expression level (geometric mean) of adult sham animals. White bars = adult sham, and dark grey bars = aged sham. Differences between groups: *p < 0.05, **p < 0.01, ***p < 0.001.

We next examined the cellular processes activated by LI-rTMS that may counter the adverse gene expression changes linked to ageing. We compared the effects of 3 daily sessions of LI-rTMS against their age-matched sham controls and observed that specific clusters of genes were modified by iTBS in *adult* cerebellum (Fig. 3A; principal component analysis – PCA turquoise), whereas BHFS altered genes mainly in the *aged* cerebellum (Fig. 3A, B, PCA dark red). The relative lack of effect of iTBS in the aged cerebellum and BHFS in the adult were confirmed by qPCR of genes regulating neuronal excitability (see Supplementary Fig. S1A). iTBS regulated only 300 genes in the adult cerebellum (Fig 3A-C). Downregulated genes were predominantly associated with developmental processes (Supplementary Fig. S2), whereas upregulated genes related to synaptic activity and ion channel function, including cation channel/transporter activity (p=0.014), calcium-related exocytosis (p=0.03), membrane repolarization (p=0.047) and synaptic modulation (p=0.0001), which cluster with glutamate secretion (p=3.33×10^-2^), synaptic plasticity (p=1.35×10^-2^) and GABAergic synaptic transmission (p=3.77×10^-2^; Fig. 3D,E).

**Fig. 3.**
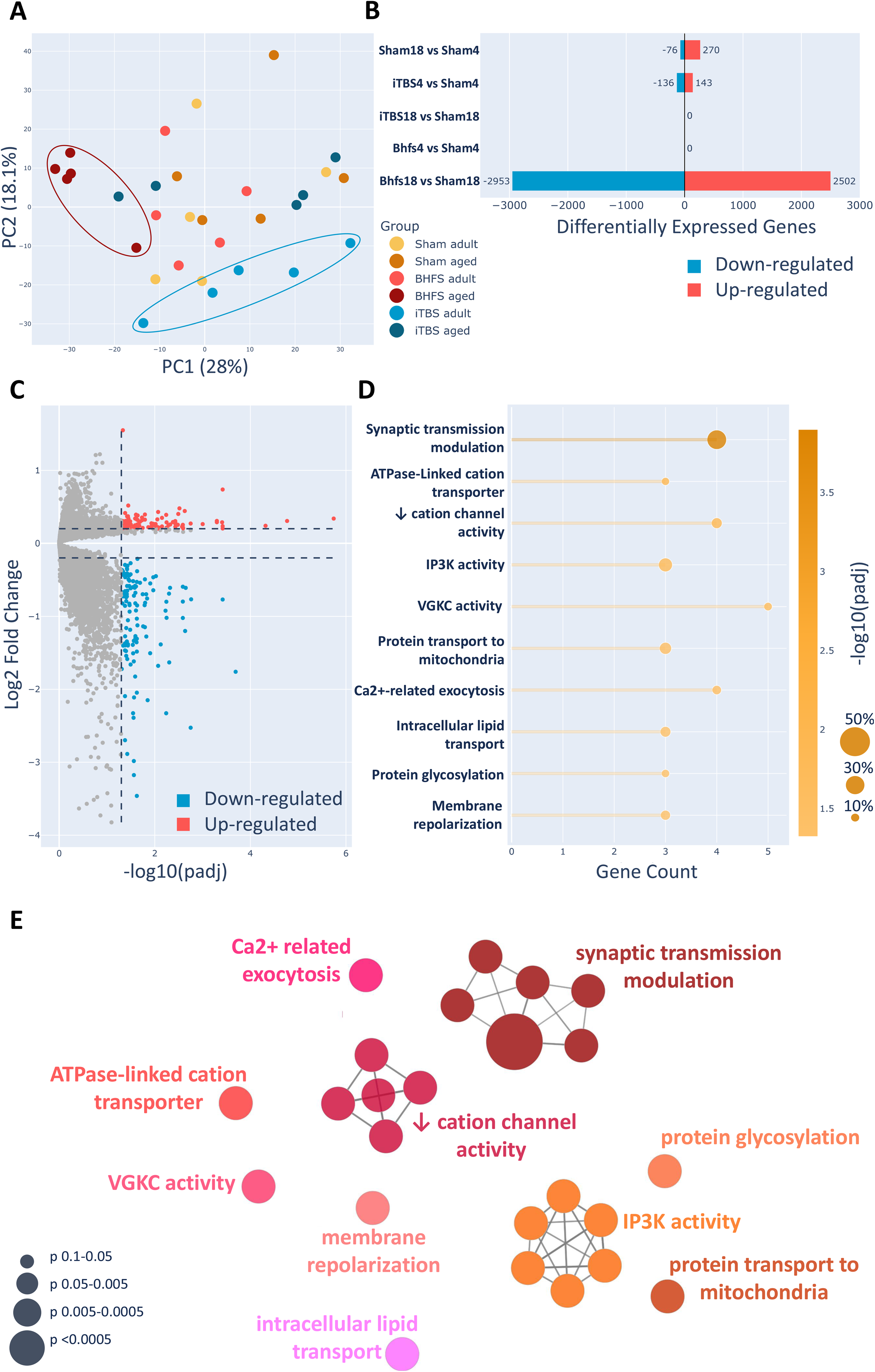
iTBS Increases Synaptic Function in the Cerebellum. **(A)** Principal Component Analysis (PCA) of RNAseq data. RNA sequencing reads were aligned using Hisat2 and quantified with FeatureCounts. Five biological replicates were analyzed per condition. Each point represents an individual sample, with colors indicating experimental groups as defined in the legend. Samples with similar gene expression profiles cluster together in the PCA plot, as indicated following iTBS to the adult, and BHFS to the aged cerebellum. **(B)** Differentially expressed genes (DEG) by condition (age and LI-rTMS treatment). Horizontal bars display the number of upregulated (red) and downregulated (blue) genes. From top to bottom, the comparisons shown are: effect of aging; iTBS in adults; iTBS in aged animals; BHFS in adults; and BHFS in aged animals. **(C)** Volcano plot of differentially expressed genes in the adult cerebellum following iTBS stimulation compared to non-stimulated controls. The vertical line indicates the significance threshold (Padj < 0.05). Genes upregulated after iTBS (Log2FC > 0.2) are shown in red, while downregulated genes (Log2FC < -0.2) appear in blue. Grey dots represent genes with non-significant or minimal expression changes. **(D)** GO showing the top biological processes (left axis) increased by iTBS. Each pathway is associated with a number of altered genes (x-axis) and an adjusted p-value gradient, reflecting enrichment significance. Bubble size indicates the proportion of upregulated genes within each biological process. **(E)** GO map showing functionally grouped networks of biological processes upregulated by iTBS. The most significant pathway (largest node) in each cluster is designated as the leading term and pathway interactions are indicated.

In contrast, the effect of BHFS was striking, altering over 5000 genes in the aged cerebellum (Fig 4 and 5) with roughly equal numbers being up- and down-regulated. Considering only those DEGs with a large fold-change (>0.4), compared to sham aged cerebellum, BHFS reduced the expression of genes involved in biological processes typically associated with ageing: peptide metabolism (p=2.93×10^-47^); oxidative stress response (p=1.47×10^-47^); inflammation (p=6.47×10^-4^); and apoptosis (p=1.21×10^-4^; Fig. 4). Conversely, it upregulated processes related to neuronal plasticity (Fig. 5), notably nervous system development (p=1.58×10^-32^), and its subgroups neuron differentiation (p=5.69×10^-19^), neurogenesis (p=4.89×10^-7^), dendrite development (p=7.59×10^-11^) and axon extension (p=7.84×10^-3^). BHFS also increased expression of genes involved in cell signalling (p=1.69×10^-10^), particularly synaptic signalling (p=2.91×10^-13^), which cluster with those regulating membrane potential (p=1.10×10^-7^), neurotransmitter release (p=1.69×10^-4^), post-synaptic organization (p=4.56×10^-8^), synaptic plasticity (p=5.64×10^-6^), and learning/memory (p=3.30×10^-3^; Fig. 5). These beneficial changes were confirmed by qPCR, with BHFS restoring the expression of several age-downregulated synaptic genes toward that of adult sham controls (Fig. 5C). Importantly, BHFS did not disturb expression in the normal adult. The only effect of BHFS in the adult cerebellum was the upregulation of *Bdnf* and *Gria1* expression (Supplementary Fig. S1B); which has been previously been demonstrated in the cerebral cortex after high intensity stimulation ^37^.

**Fig. 4.**
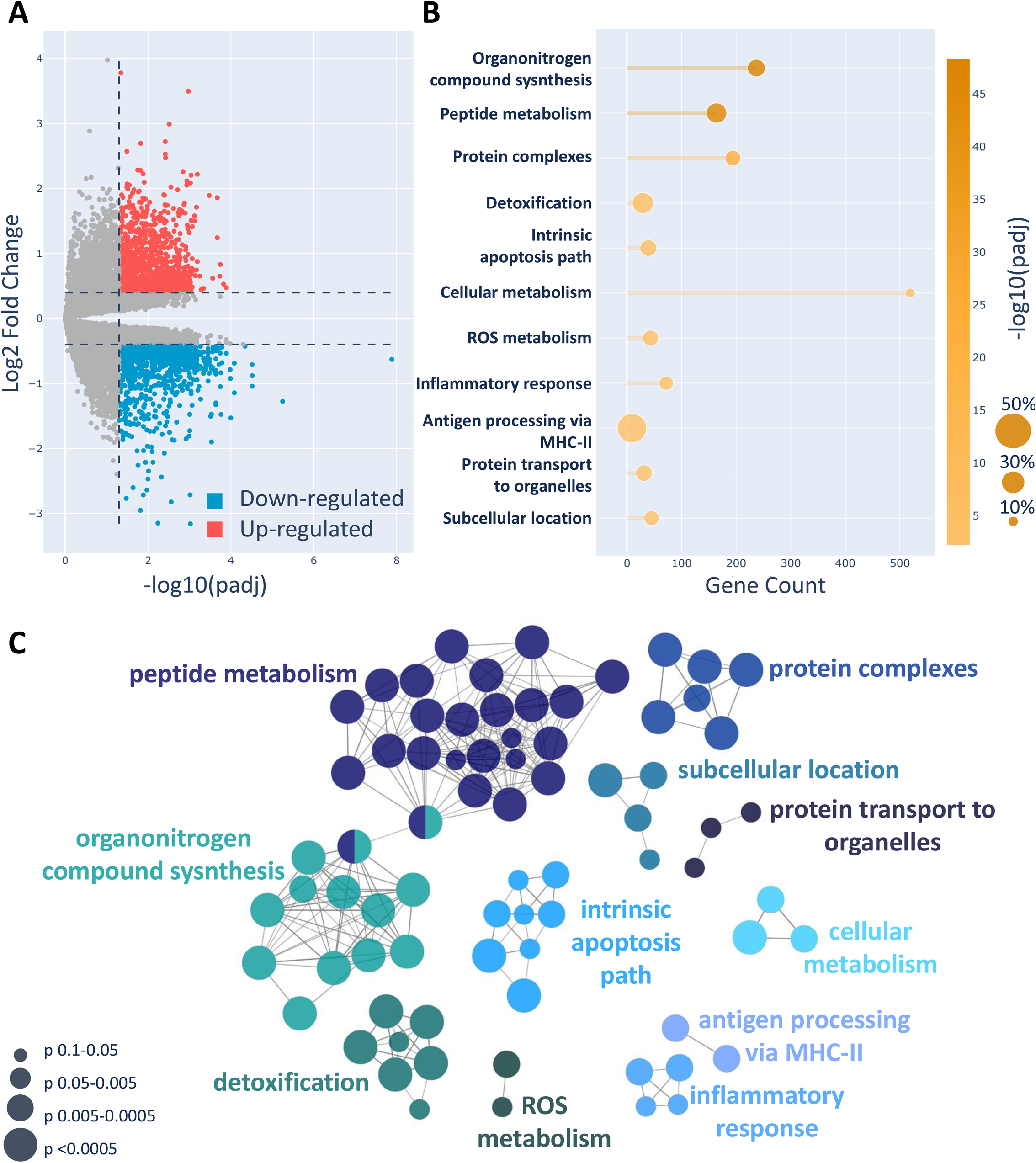
BHFS Decreases Ageing-Related Biological Processes in the Cerebellum. **(A)** Volcano plot of differentially expressed genes in the aged cerebellum after BHFS stimulation compared to non-stimulated controls. The vertical line indicates the significance threshold (Padj < 0.05). Genes upregulated following BHFS (Log2FC > 0.4) are shown in red, while downregulated genes (Log2FC < -0.4) are shown in blue. Grey dots represent genes with non-significant or minimal expression changes. **(B)** GO of the main biological processes (left axis) reduced by BHFS. Each pathway is associated with a number of altered genes (x-axis) and an adjusted p-value, reflecting enrichment significance. Bubble size indicates the proportion of downregulated genes within each biological process. **(C)** GO map of functionally grouped networks of biological processes downregulated by BHFS. The most significant pathway (largest node) in each cluster is designated as the leading term and pathway interactions are indicated.

**Fig. 5.**
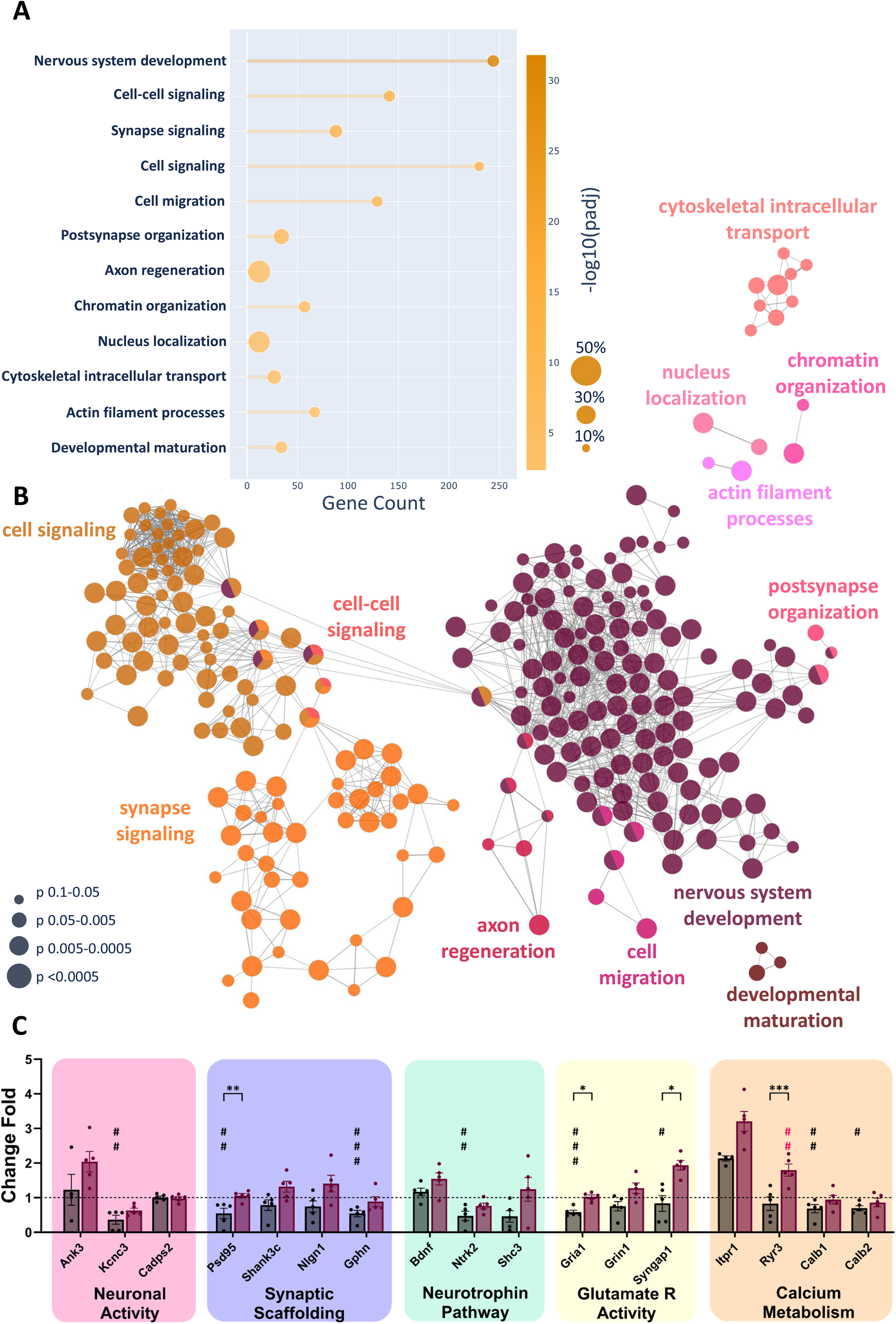
BHFS reverses age-related decline in expression of cerebellar neuroplasticity genes. **(A)** GO of biological processes increased in the cerebellum by BHFS showing the number of genes (x-axis) in the main upregulated processes (left axis), their significance (adjusted p-value gradient; right axis) and the proportion of genes that are changed. **(B)** GO map showing a functionally-grouped network of biological processes that are upregulated by BHFS. The most significant pathway (largest node) in each cluster is designated as the leading term. Pathway interactions are indicated. **(C)** qPCR of some cerebellar neuroplasticity genes. Data are expressed as relative expression normalized to the adult sham group, according to the Pfaffl equation. The horizontal dotted line represents the mean expression level (geometric mean) of adult sham animals. Dark grey bars indicate expression in aged sham, and violet bars represent aged BHFS group. Differences between groups: *p < 0.05, **p < 0.01, ***p < 0.001, ****p < 0.0001. Comparisons to adult sham (data not on the graph): # p < 0.05, ## p < 0.01, ### p < 0.001.

### 3 daily sessions of LI-rTMS increases PC spine density and alters spine morphology

To understand how these gene expression changes may contribute to LI-rTMS-induced repair of age-related cerebellar decline, we examined Purkinje cell morphology after 3 days of 10 minutes BHFS, as this was the stimulation protocol inducing an extensive range of DEGs in the aged cerebellum.

Purkinje cells from aged mice had a lower dendritic spine density than those from adults (2-way ANOVA F_1,80=_ 11.82, p=0.0009), and LI-rTMS increased spine density in both groups (adult, p<0.0001; aged, p=0.026; Fig 6A). This increase was paralleled by spine subtype changes: stubby-spine density was increased in both adult and aged Purkinje cells (2-way ANOVA, F_1,80=_ 9.09 p=0.0034; effect of stimulation F_1,80=_ 36.52 p<0.0001), while thin/mushroom-spines were only increased by LI-rTMS in adults (2-way ANOVA, F_1,80=_ 7.78 p=0.0066; effect of stimulation F_1,80=_ 9.38 p=0.003; Fig 6B-D). The increase in total spine density induced by LI-rTMS is thus primarily due to a greater increase in stubby-spines than in thin/mushroom spines. Additionally, LI-rTMS modified the morphology of the thin/mushroom spine population, reducing their head diameter (Kolmogorov-Smirnoff test; p=0.002; Fig 6Ei) and length (Kolmogorov-Smirnoff test; p<0.0001; Fig 6Eii) in adult animals, but only their length in aged animals (Kolmogorov-Smirnoff test; p=0.003; Fig 6Eiv). These morphological changes suggest that BHFS LI-rTMS generated more newly-formed (i.e., stubby) spines, which is consistent with a plasticity state ^38^. Despite these robust dendritic spine changes, 3 sessions of LI-rTMS did not alter dendritic tree size or complexity (Supplementary Fig. S3).

**Fig. 6.**
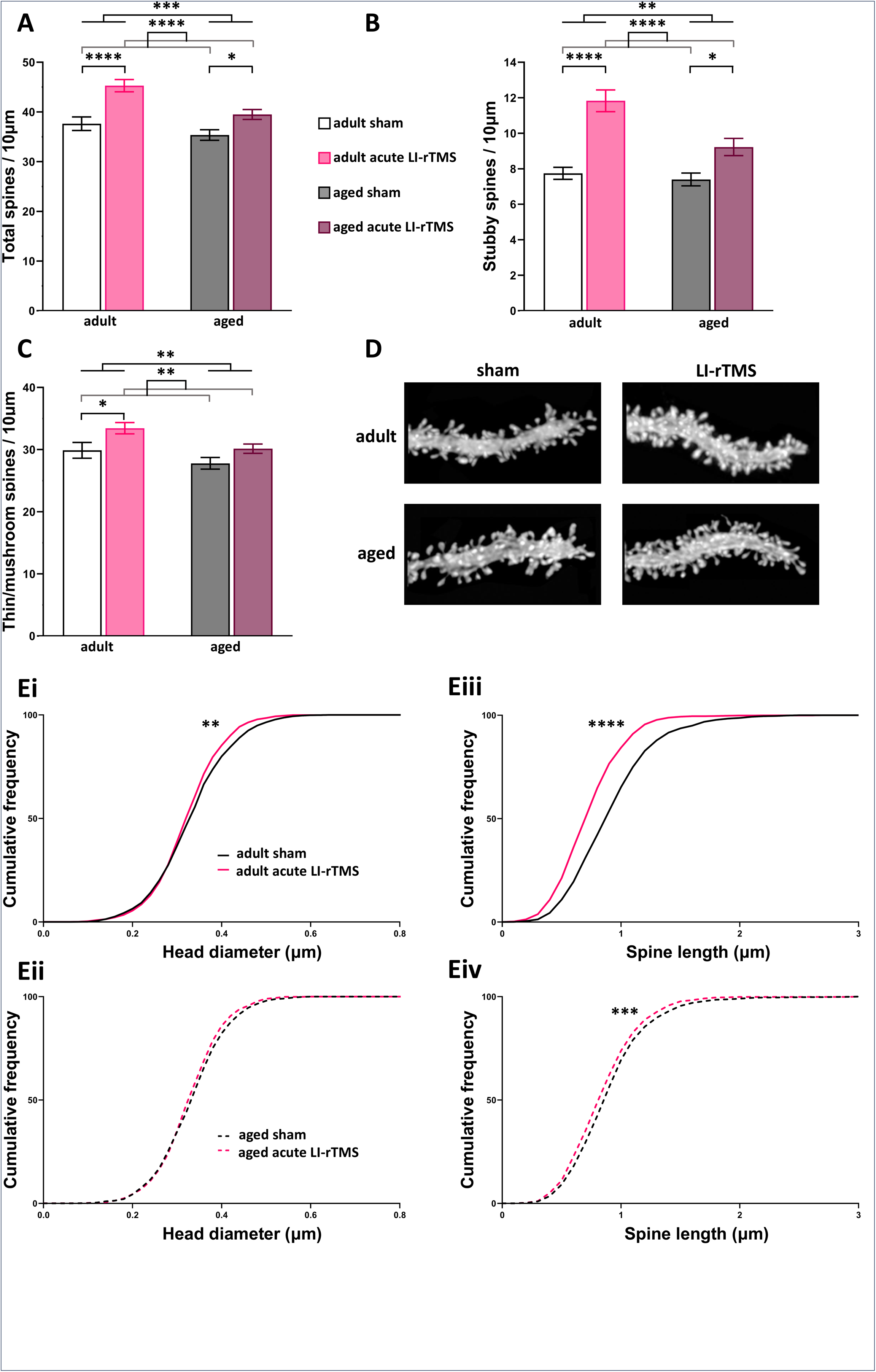
Acute LI-rTMS (3 days) increases Purkinje cell dendritic spine density and modifies spine morphology in both adult and aged mice. **(A)** Aged animals have significantly lower spine density than adult animals (Two-way ANOVA F_1,80=_11.82 p=0.0009; stimulation F _1,80 =_ 25,17, p<0,0001), which was increased by LI-rTMS in both adult (p<0.0001) and aged mice (p=0.026; *n*=5, N=18-25). **(B)** Stubby spine density in aged animals was also lower than in adults (Two-way ANOVA F_1,80=_9.095 p=0.0034; stimulation F_1,80 =_ 36,52, p<0,0001) but was increased by LI-rTMS (adult p<0.0001; aged p=0.017). **(C)** Thin/mushroom spine density was also lower in aged vs. adult animals (Two-way ANOVA F_1,80=_7.782 p=0.0066; stimulation F_1,80 =_ 9,38 p=0,003), and LI-rTMS increased this density only in adults (p=0.025). **(D)** Example of PC tertiary dendrites in adult and aged mice, illustrating the increased density of dendritic spines induced by LI-rTMS (right) compared to sham (left). **(E)** Acute LI-rTMS alters morphology of thin/mushroom spines in adult and aged mice (*n*=5; *N*=1575-1896). **Ei** - head diameter is reduced by LI-rTMS in adults (Kolmogorov-Smirnoff test; p=0.002) but not in aged animals (**Eii** p>0.05). However, stimulation decreased thin/mushroom spine length compared to sham, in both age groups (Kolmogorov-Smirnoff test; **Eiii** p≤0.0001 and **Eiv** p=0.003, respectively). Intergroup differences: *p≤0.05; **p≤0.01; ***p≤0.001; ****p≤0.0001. *n* = Number of animals; *N*= Number of tertiary dendrites analysed.

### Long-term LI-rTMS alters Purkinje Cell dendritic morphology

Given the effects of just 3 days of 10 minutes per day LI-rTMS, we applied the same stimulation over 4 weeks, to assess whether long-term cerebellar LI-rTMS modifies ageing Purkinje neuron morphology and improves age-related behavioural deficits.

Surprisingly aged Purkinje cells have a larger dendritic area than adult Purkinje cells (2-way ANOVA F_1,123=_731.8, p<0.0001; Fig 7 D). Moreover, 4 weeks LI-rTMS greatly increased the Purkinje cell dendritic density in both age groups (2-way ANOVA F_1,123=_731.8, p<0.0001; stimulation F_1,123=_414.4, p<0.0001; Fig 7D), but not height or width (Fig. 7B-C). Sholl analysis showed that this was due to increased dendritic branching in the middle section of the dendritic tree, particularly in the adult LI-rTMS treated Purkinje neurons (2-way ANOVAs with Bonferroni post-hocs all p<0.02; Fig7E).

**Fig. 7.**
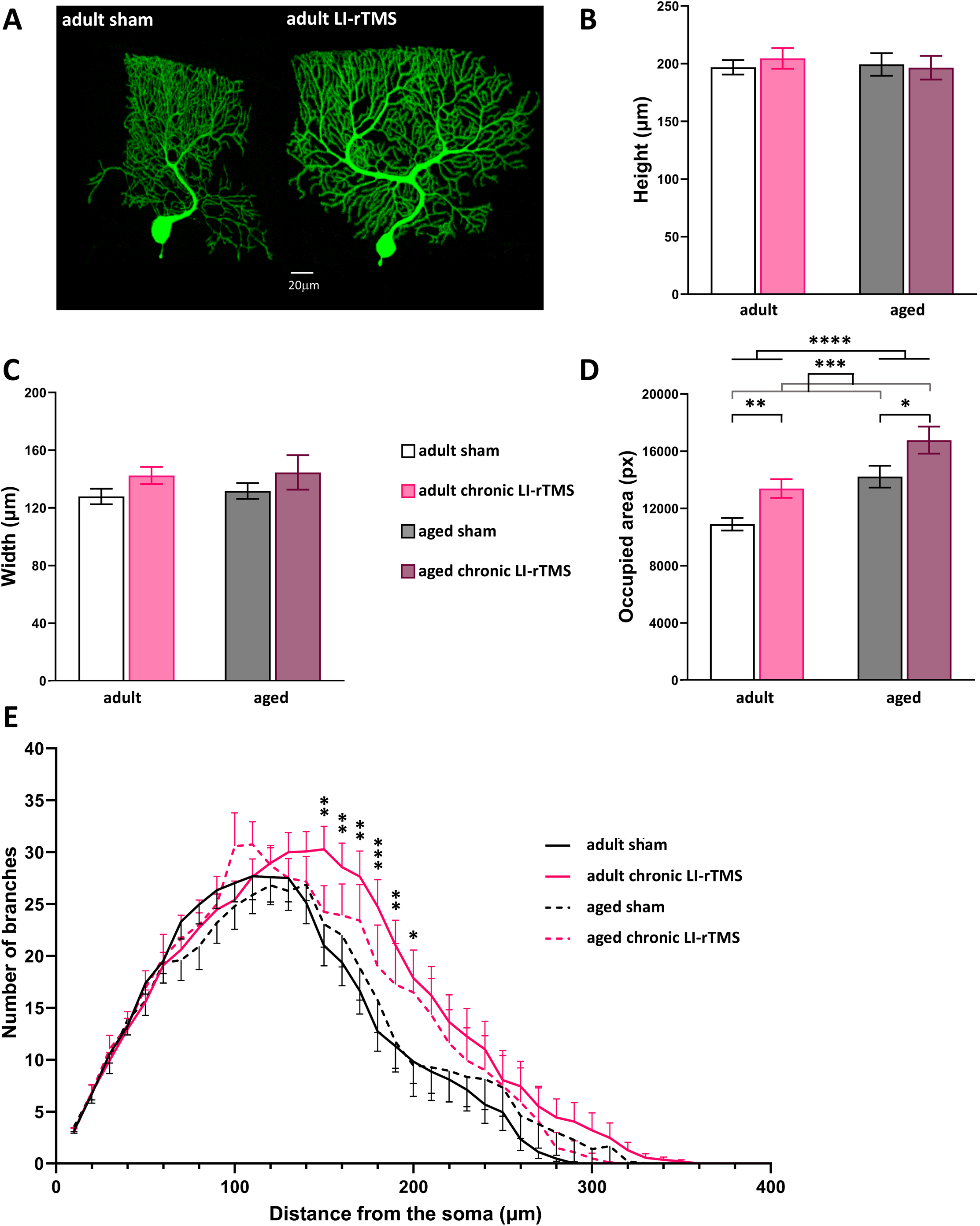
Long term LI-rTMS (4 weeks) increases Purkinje cell dendritic density and complexity. **(A)** Example of PCs showing the increased dendritic tree size induced by LI-rTMS in adult mice (right) compared to sham stimulation (left). **(B)** Dendritic height was the same at both ages (Two-way ANOVA F_1,123=_0.1008 p=0.7514; stimulation F_1,123 =_ 0.0748 p=0.7849), with no effect of 4 weeks LI-rTMS (Bonferroni post –hoc p>0.05). Adult sham *n*=16, *N*=49; adult LI-rTMS *n*=15, *N*=33; aged sham *n*=9, *N*= 25; aged LI-rTMS *n*=10, *N*=20. **(C)** Chronic LI-rTMS did not increase dendritic tree width at either age (Two-way ANOVA F_1,123=_0.1730 p=0.6782; stimulation F_1,123 =_ 3.69 p=0.057; Bonferroni posthocs both adult and aged p>0.05) **(D)** LI-rTMS increased dendritic density compared to sham (Two-way ANOVA F_1,123=_24.71 p<0.0001; stimulation F_1, 123 =_ 13.99 p=0.0003) in both adult (p=0.0051) and aged mice (p=0.039). **(E)** Sholl analysis, quantifying the number of dendritic branch points as function of distance from the soma, shows that long-term LI-rTMS (4 weeks) increases PC dendritic tree complexity in its middle section (150-200µm from the soma) in adult animals (Two-way ANOVA and pairwise comparisons, Bonferroni correction, all p≤ 0.02). Intergroup comparisons: *p≤0.05; **p≤0.01; ***p≤0.001; ****p≤0.0001. *n*= number of animals; *N*= number of Purkinje cells

In addition, although LI-rTMS did not change spine density on either adult or aged Purkinje cell dendrites after this longer protocol, it did alter spine morphology (Fig. 8). Compared to adult mice, thin/mushroom spines of aged mice had larger head diameters. LI-rTMS reduced head diameter of these spines (Kolmogorov-Smirnoff test) in both adult (p=0.0007; Fig. 8Biii) and, notably, aged mice (p<0.0001; Fig. 8Bii). Thus, the enlarged spine heads on aged Purkinje dendrites returned toward the normal adult size (p=0.02; Fig. 8Biv).

**Fig. 8.**
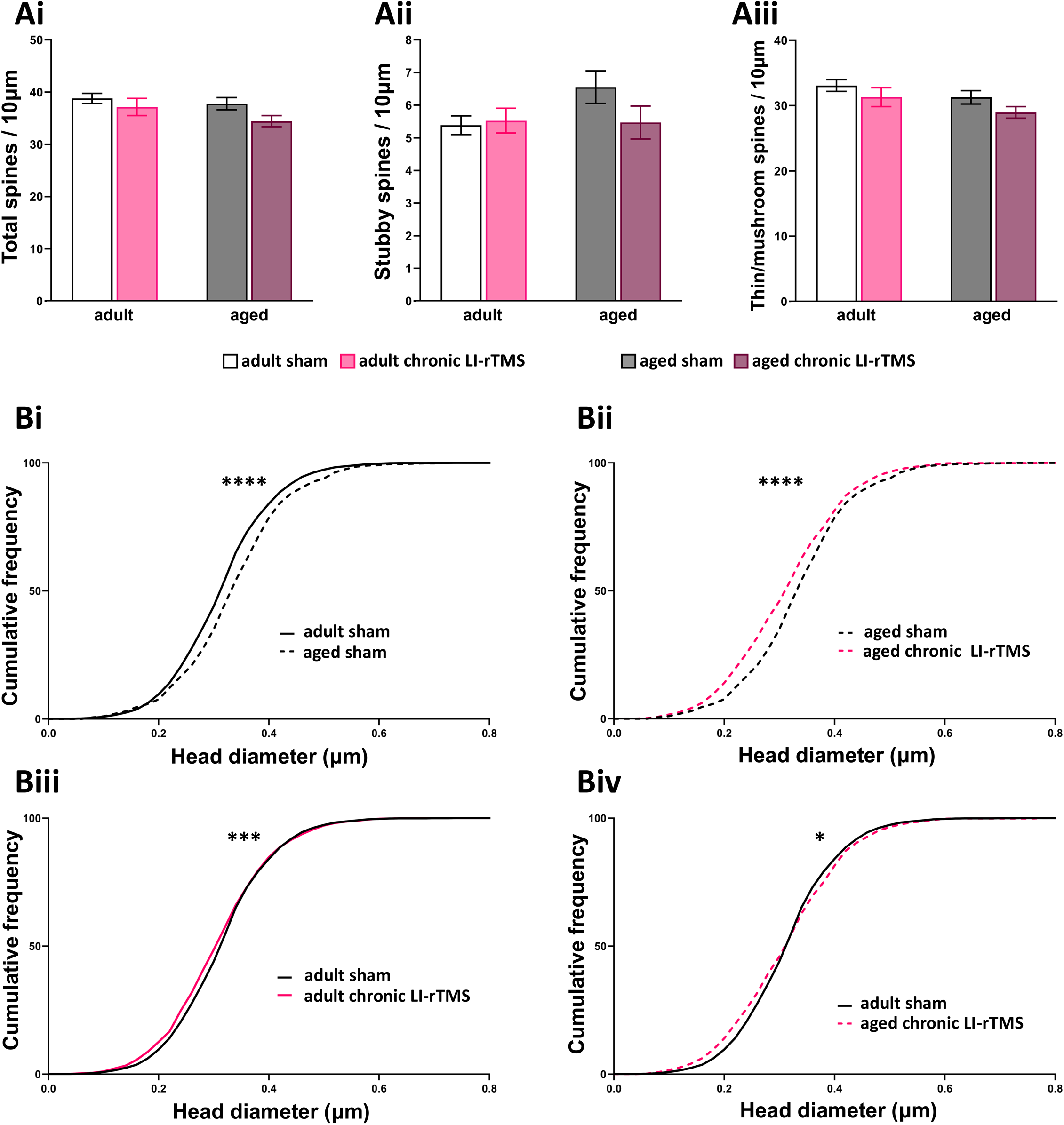
Long term LI-rTMS (4 weeks) does not change spine density but modifies spine morphology in both adult and aged animals. **(A)** There was no difference between groups in the density of all, stubby or thin/mushroom spines (Two-way ANOVA; all p>0.05; adult sham *n*=16, *N*=40; adult LI-rTMS *n*=10, *N*=35; aged sham *n*=9, *N*= 20; aged LI-rTMS *n*=10, *N*=18. **(B)** Purkinje cells from aged sham mice have larger thin/mushroom spine heads than adult sham Purkinje cells (**Bi**: Kolmogorov-Smirnoff (K-S) distribution, p<0.0001). LI-rTMS decreases spine head diameter in adult (**Biii**: K-S distribution, p=0.0007) and particularly in aged (**Bii**; K-S distribution p≤0.0001) mice, so that head diameters of thin/mushroom spines on aged Purkinje cells approximate to normal adult spine head diameter (**Biv**: K-S distribution, p=0.02). Adult sham *N*=3084, adult LI-rTMS *N*=3045; aged sham *N*=915; aged LI-rTMS *N*=707. *p≤0.05; **p≤0.01; ***p≤0.001; ****p≤0.0001 *n*= number of animals; *N*= number of cells (A) or spines (B)

These data indicate that ongoing LI-rTMS continues to modulate Purkinje cells, increasing dendritic branching, and therefore total spine numbers, as well as favoring more plastic thin/mushroom spines (smaller head diameter).

### 4-weeks LI-rTMS improves spatial memory

We have shown that Purkinje dendritic spines are rapidly induced by LI-rTMS and that ongoing stimulation increases dendritic tree branching (Fig. 7). These modifications to Purkinje cell connectivity could potentially optimise function of this circuitry to improve associated behaviours: motor coordination and spatial learning and memory ^14,15,39^.

On the accelerating rotarod, all groups improved their performance, i.e. motor coordination, during the 5-day learning period (Friedman nonparametric RMANOVA and Wilcoxon ranked test, P=0.0014; Fig. 9A), with no consistent differences between LI-rTMS and sham treated groups. A more discriminating test of cerebellar function is spatial learning and memory using the Morris Water Maze ^39^. To ensure all animals could orientate themselves and swim to a platform, mice performed 2 trials when the platform was visible. There were no significant differences between any groups for swim speed, distance travelled or the latency to the platform (Kruskal-Wallis test, p>0.05; Supplementary Fig. S4).

**Fig. 9.**
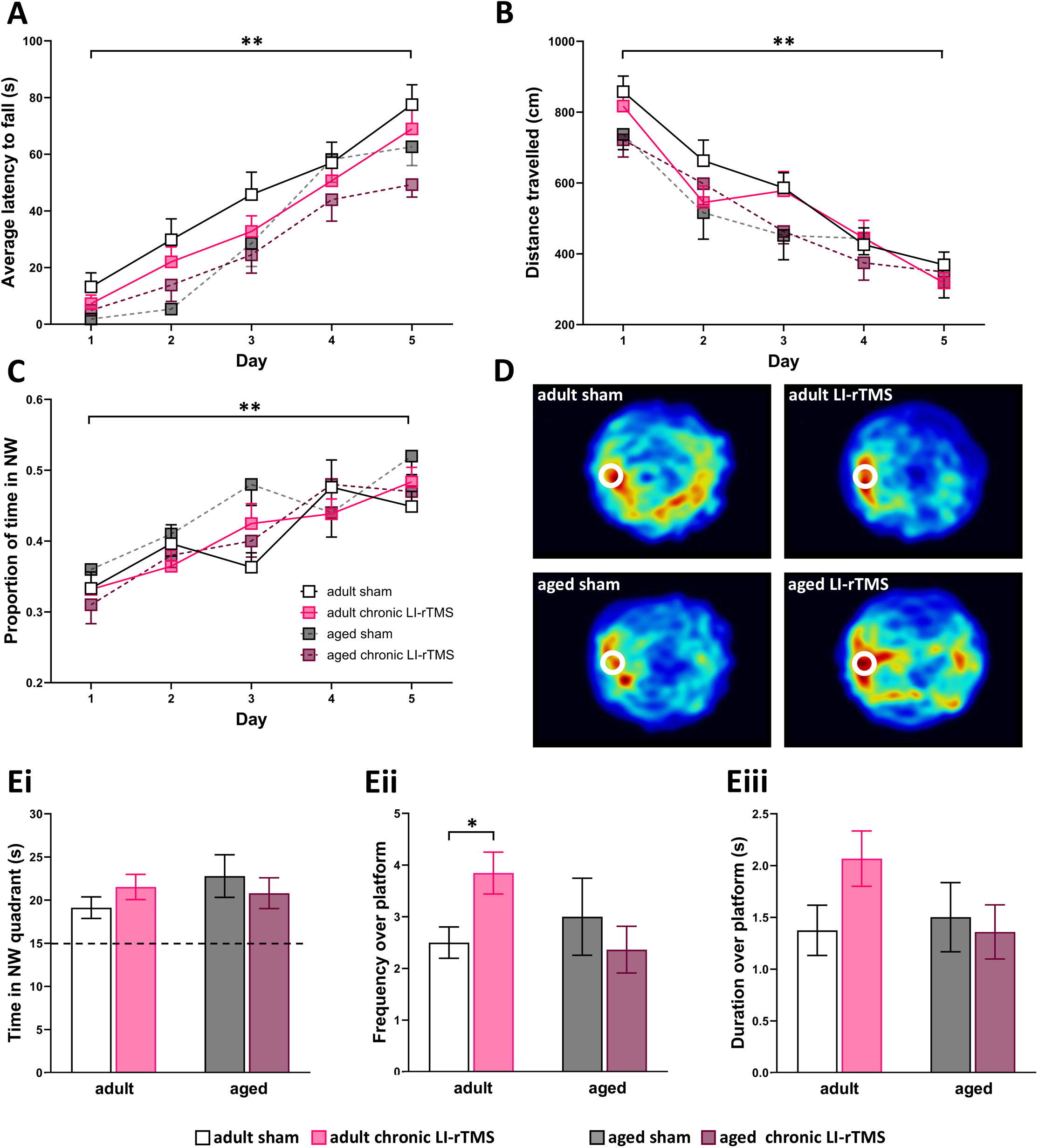
Cerebellar LI-rTMS does not alter motor coordination or spatial learning but improves spatial memory. **(A)** All groups learned the accelerating rotarod task during 5 days (Friedman non-parametric RMANOVA, day 1 vs day 5 p<0.0014). No differences were observed between groups (Kruskal Wallis all p>0.05). **(B, C)** In the learning phase of the Morris Water Maze (MWM) all groups improved their performance as measured by shorter distance travelled **(B)** and greater time spent in the correct quadrant **(C)** (Friedman RMANOVA; day 1 vs day 5 p<0.0014 and p<0.007 respectively). No differences were observed between groups (Kruskal Wallis all p>0.05). **(D)** Heat map representing the merged localisation of mice during the probe test of the Morris Water Maze. Adult animals treated with LI-rTMS spent more time over the platform location (white circle) than adult sham treated animals, as indicated by the red colour. This effect was not observed in aged mice. **(Ei)** In the probe test of the MWM, all mice spent more time than chance (15s) in the target quadrant, indicating learning. (**Eii)** LI-rTMS increased the number of times adult mice crossed the platform location (2-way ANOVA and Bonferroni’s post-hoc p=0.041) and (**Eiii)** the duration spent over it. This effect was not seen in the aged animals (p>0.05).

During the 5 days learning to find the hidden platform, all mice improved their performance (RMANOVA), reducing the distance travelled (p<0.03; Fig. 9B) and latency to reach the platform (p<0.001), while increasing time spent in the correct quadrant (p<0.02; Fig. 9C), but without a clear effect of LI-rTMS (Fig. 9B-D). However, the real test of this learning comes through the probe test, in which mice demonstrate memory by swimming to the platform’s location even when the platform has been removed (Fig. 9E). While mice in all groups learned the platform’s location (Fig. 9Ei), LI-rTMS improved memory of the platform’s location in adult mice (2-way ANOVA Bonferroni’s post-hoc p=0.041) with increased number of crossings and time spent over the platform’s previous location (Fig. 9Eii-iii).

## DISCUSSION

We have investigated the effects of LI-rTMS on the cerebellum, a brain region that undergoes degenerative changes early in ageing and which modulates cerebral neuronal networks and associated cognitive and emotive function ^16^. We observed that LI-rTMS rapidly modifies gene expression in a stimulation-dependent manner, reducing adverse cellular processes that underlie ageing and increasing those implicated in synaptic plasticity. In parallel, LI-rTMS increased spine density on dendrites of the central cerebellar neuron, the Purkinje cell. Purkinje cell changes were consolidated by 4 weeks of daily LI-rTMS to develop a larger, more complex dendritic tree. All groups showed similar function in cerebellar-related tasks, although LI-rTMS improved spatial memory in adult mice.

### LI-rTMS rapidly activates stimulation- and age-specific cerebellar responses

LI-rTMS modified cerebellar gene expression in stimulation- and age-dependent manners. iTBS altered gene expression (∼300) mainly in the normal adult cerebellum, whereas BHFS modified an extensive number of genes (>5000) in aged tissue (Fig. 1).

The change in gene expression after iTBS in the normal adult cerebellum were primarily upregulation of genes involved in neuronal excitability and synaptic function and plasticity, consistent with the known excitatory effect of iTBS stimulation in humans ^40^, and reduced inhibitory interneuron firing ^41^. However, it contrasts with the effect of medium-intensity (120 mT) TBS on the cerebral cortex (both motor and somatosensory) in which the few DEGs were mainly downregulated: ribosomal functions by iTBS and synaptic genes by cTBS ^8,11^. These different effects between similar stimulation protocols applied to different brain regions confirm the need for fundamental studies to identify what rTMS does to different brain regions *per se*, as well as to their down-stream targets. Moreover, the very great difference in transcriptomic modifications induced in aged vs. adult cerebellum confirms previous findings that a circuit’s functional state modifies the way it responds to rTMS, with normal neural circuits being much less reactive than dysfunctional ones ^36,42^.

Gene expression changes were associated with modifications in dendritic spines. In both adult and aged animals 3 days of LI-rTMS increased spine density, particularly of stubby spines, consistent with a state of morphological and synaptic plasticity ^38^. Spine remodelling continues in the adult brain ^38,43^ underlying memory formation ^44,45^ and adaptive brain changes to environmental input ^46^. Here we show that LI-rTMS can induce spinogenesis and morphological spine plasticity even in the aged brain. Our data show that in aged Purkinje cells thin/mushroom spines have unusually large head diameter (Figs. 6 and 8). This morphology is usually indicative of stable spines ^47^, which is consistent with less synaptic flexibility. LI-rTMS however, reduced these enlarged head diameters almost to the normal adult size, and this was sustained for at least 2 weeks after the end of stimulation, indicating that LI-rTMS had restored age-impaired synaptic plasticity. Moreover, since LI-rTMS does not necessarily elicit neuronal firing ^48^, this stimulation appears to induce spine changes without action potentials. This is of particular interest because impaired activity-dependent spine morphogenesis underlies many neurological disorders, not just ageing ^38,49^.

These acute effects of LI-rTMS on spine structural plasticity are present in both adult and aged animals, but were more pronounced in the adult group, indicating a lower level of structural synaptic plasticity induced by LI-rTMS with age. Poorer acute responses to rTMS of aged people have been previously demonstrated ^7,50,51,52^ as well as reduced spine plasticity in the aged motor cortex in response to iTBS ^53^. However, although aged neurons are less responsive electrophysiologically ^54,55^, consistent with reduced neuronal plasticity during ageing ^52,56^, our gene expression data show that aged cerebellar neurons are still able to respond dramatically when appropriately stimulated, creating a huge potential for facilitating neuronal function.

Our results indicate that as little as 3 days of 10minutes per day cerebellar LI-rTMS is able to modify neuronal gene expression and increase the number of synaptic sites on Purkinje cells, reshaping connectivity within aged cerebellar neuronal circuits and therefore potentially restoring function in the aged cerebello-cerebral network.

### Chronic LI-rTMS changes both Purkinje cell dendritic morphology and spatial memory

A key finding of this study is that the promising initial effects of LI-rTMS to the cerebellum are consolidated with longer treatment, increasing Purkinje cell dendritic density and complexity in both adult *and* aged animals. It is tempting to speculate that the rapid increase in spine density after 3 days of treatment is confirmed with their accommodation onto the newly grown dendrites of the expanded dendritic tree.

Increases in spine density and Purkinje cell dendritic tree size induced by LI-rTMS are consistent with morphological changes observed in Purkinje neurons following periods of motor learning or environmental enrichment ^57,58,59^ which modulate synaptic efficacy ^60^ Indeed, dendritic branching complexity influences the efficacy of action potential propagation along dendrites ^61^ and the computational power of the neuron ^62^ which in turn impact upon integration of synaptic input and induction of plasticity ^62,63^.

In addition, spatial memory was improved in the LI-rTMS-treated adult animals, suggesting a correlation between morphological and behavioural effects. Although our stimulation targeted the cerebellum, and the cerebellum regulates the egocentric component of spatial learning and memory ^39,64,65^, it is possible that stimulation overspill to the hippocampus and parietal cortex, even at a much reduced intensity (estimated < 1 mT), may have contributed to improved spatial memory performance. In contrast, behavioural improvement was not seen in aged animals, despite Purkinje cell morphological changes equivalent to those in adults, suggesting that improved cerebellar function in these animals is not reaching other brain regions, possibly because of age-related atrophy of the cerebello-efferent axons (superior cerebellar peduncle; ^14^).

### Conclusions

These studies show that LI-rTMS, which induces an electric field too weak to directly elicit action potentials in the underlying neurons, nonetheless changes rapidly gene expression after only a few stimulation sessions. These effects could be induced in both the young adult and aged cerebellum, but through very different stimulation protocols. LI-rTMS also modifies spine density and morphology, and with longer treatment, dendritic complexity, thus changing cerebellar neural circuit structure. Significantly our data show that appropriate rTMS protocols can create structural change in neurons that are highly vulnerable to ageing, returning both gene expression and spine morphology to normal adult parameters. This suggests that treating vulnerable brain regions early during ageing, in order to restore normal circuit connectivity and repair age-related cognitive decline, is a highly promising treatment for our increasingly aged population.

## MATERIALS AND METHODS

### Animals

All animal procedures were carried out under French ethics guidelines and were approved by the local ethics committee (“Comité Charles Darwin, N° 5 »). C57Bl/6J wild-type mice were bred at the Institut de la Longévité, Ivry-sur-Seine, France. They were housed in single-sex cages on a 12 h light-dark cycle and with water and food *ad libitum*.

Mice aged 4-5 months (adult) and 16-17 months (aged) received LI-rTMS for 10 minutes per day for 3 days. Following the last stimulation, animals were euthanized and the cerebellum taken at 5 hours for RNA sequencing or at 20h for Purkinje cell visualization. A second cohort of adult and aged mice were treated with LI-rTMS for 4 weeks, followed by 2 weeks of behavioural testing and subsequent visualization of Purkinje cells.

### Magnetic Stimulation

Low intensity rTMS was applied to the cerebellum as previously described ^27,34,36^. Focal LI-rTMS used a custom magnetic coil designed for mice (copper wire, 300 windings, 16Ωm, inner diameter 5mm, outer diameter 8mm), and powered by an electromagnetic pulse generator (EC10701; Global Energy Medicine Pty Ltd, Perth, Western Australia). The frequency used for stimulation was either BHFS (62.5ms trains of 20 pulses repeated at 9.75 Hz) or iTBS as previously described ^27^. The intensity of the magnetic field was 9mT at the tissue, as measured with a Hall device.

For stimulation, each animal was placed in a small box to limit mobility and the LI-rTMS stimulation coil held above the posterior scalp, over the cerebellum, for 10 minutes each day. Mice showed no signs of discomfort or stress during or after the stimulation. Control mice received sham stimulation, with the stimulator turned off. The sound emitted by the coil is below the audible threshold of mice ^36^ and coil temperature does not change during stimulation ^34^

### Behavioural analysis

#### Rotarod - motor coordination

Mice were placed on a rotating rod and the latency to fall was measured. We used an accelerating rotarod (TSE Systems, Germany) with a 3 cm diameter. Initial speed was 4 rpm; frequency increased by 4 rpm every 30 seconds for a maximum of 5 minutes (40rpm). Mice received 3 trials per day for 5 days.

#### Morris Water Maze - spatial learning and memory

##### Learning

Each mouse was placed in a 120 cm circular tank filled with water (23°C) and were trained to use visible external cues to find an invisible clear Plexiglas platform (10cm diameter) hidden 1 cm under the water’s surface. Training took 6 trials per day for 5 consecutive days, with the platform always in the northwest quadrant and the mouse release from one of 4 randomly-selected starting points (north, east, south or west). Mice were given 60 seconds to locate and climb onto the platform. If the mouse did not succeed, it was placed on the platform and held there for 15 seconds. Mice were tracked using with EthoVision XT (Noldus). Parameters measured were latency to platform, path length, time spent in the northwest quadrant, swim velocity and the percentage of direct swims, for which daily averages were calculated for each mouse.

##### Probe test

Two hours following the final trial., the platform was removed and the mouse released from the south or east positions. Parameters measured were the latency to the platform location, frequency of crossings and duration over the it, time in the northwest quadrant, mean distance from the platform position and number of direct swims.

##### Visible Platform

On the final day, the platform was moved to the south east quadrant and was raised above the surface with a highly visible object and light marking its. Parameters measured were distance moved, latency to platform and swim velocity.

### Purkinje cell morphology

Sagittal cerebellar slices were prepared from LI-rTMS- and sham-treated animals. Mice were sacrificed by rapid decapitation and the cerebellum removed. Sagittal vibratome slices (250 μm) were cut at 4°C in an oxygenated external solution containing (in mM) 125 NaCl, 2.5 KCl, 2 CaCl2, 1 MgCl2, 25 NaHCO3, 1.25 NaH2PO4, 25 Glucose). In midline slices, Purkinje cells in lobule VI/VII were patch-clamped using an internal solution including 5 mM biocytin. After filling several PCs, slices were fixed in 4% paraformaldehyde overnight at 4°C. To visualize the biocytin-filled cells, slices were washed in phosphate buffered saline (PBS) containing 0.25% Triton and incubated with Streptavidin AF488 conjugate (1:400; Invitrogen, Molecular Probes) diluted in PBS containing 0.25% Triton X, 0.2% gelatine, and 0.1M lysine. Following washes with PBS-T 0.25%, slices were mounted in Mowiol (Sigma).

Confocal (Leica TCS SP5, Leica) Purkinje cell image stacks (0.5μm step) were acquired at 40x (NA=1.25) or 63x (NA=1.4) objective, with an argon 488nm laser. Image-stacks were flattened using maximum pixel intensity, and the resulting image was analysed. Measurements were made of the dendritic tree: total area, height (soma to the arbor edge at the pial surface), width at the apex, base and mid-height, from which an average width was calculated. The image was then thresholded, and the fraction of the arbor area containing labelling indicated the area occupied by dendritic branchlets. Dendritic branching complexity was analysed by the Sholl method. The ImageJ plugin tubeness (ơ = 0.8; http://rsb.info.nih.gov/ij/index.html) was applied to the z-stack, then images were flattened for Sholl analysis using concentric spheres at 10µm intervals ^66^.

Spine density and morphology was analysed on confocal image stacks (63x objective with 4x optical zoom, step 0.13μm) of two distal Purkinje dendrites (3-5 PCs per animal). Images were deconvoluted using Huygens 3.7 software (Scientific Volume Imaging). For each neuron, a tertiary distal dendritic segment of 20-40 µm in length was analysed with Neuronstudio (^67^ http://research.mssm.edu/cnic/tools.html_),_ which detected spines in thin, stubby or mushroom categories. Manual correction was required for a minority of spines. The total spines, and number within each category, was counted for each cell and expressed as the density of spines per 10µm of dendrite. Spine head diameter and length were automatically calculated and data imported into Graphpad Prism to create frequency distribution of spine head diameter and length for each group ^68^.

### Gene expression and RNA-sequencing

5 hours after the third LI-rTMS session to adult and aged mice, mice were euthanized and the cerebellum harvested. Total RNA was extracted using TRIzol (Ambion) according to the manufacturer’s instructions and stored at −80°C.

For qPCR, complementary DNA (cDNA) was transcribed from 500 ng of total RNA using the High-Capacity cDNA Reverse Transcripvon Kit (Applied Biosystems) and amplified with a CFX Opus Real-Time PCR System (Bio-Rad Laboratories, USA). Results were analysed on the Bio-Rad CFX Maestro soxware (Bio-Rad Laboratories, USA) according to Pfaff’s equavon using *Hprt* and *Mrpl32* as housekeeping genes. Primer sequences are in Supplementary Table 2. Normalized mean expression levels were used to determine differentially expressed genes between each group.

For RNA sequencing, RNA quality was assessed using Agilent High Sensitivity RNA ScreenTape System. 300ng RNA, with an RNA Integrity Number (RIN) >8, were used for subsequent sequencing (Supplementary Fig. S5 and Supplementary Table 1 for workflow).

### Statistical analysis

Data were tested for normal distribution and homogeneity of variance; square-root or log^e^ transformation was applied if needed. When appropriate, two-way ANOVA and Bonferroni post hoc tests were used, with treatment and age as between-subject factors. If interactions between factors were observed, one-way ANOVA and pairwise comparisons with Tukey post hoc tests were performed. Non-normal data were analysed by Kruskal-Wallis with Dunn-Bonferroni correction or Mann-Whitney U tests.

For the learning phase of Morris Water Maze, two-way ANOVA repeated measures tests were done. Rotarod data were analysed by Friedman non-parametric test and Wilcoxon ranked post-hocs. For spine morphology analysis Kolmogorov-Smirnoff distributions were compared. RNA sequencing data were analysed by Gene Ontology (GO) of differentially expressed genes (DEGs; Supplementary Fig. 5 and Supplementary Table 1 for workflow) and qPCR by ANOVA. Statistical analyses were performed using GraphPad Prism v10, SPSS v22 or R software and α set at 0.05.

## FIG. LEGENDS

**Supplementary Fig. 1. LI-rTMS has different effects depending on age and pattern.**

Gene expression data are expressed as fold change or relative expression normalized to the adult sham group, according to the Pfaffl equation. The horizontal dotted line represents the mean expression (geometric mean) of adult sham animals. Dark grey bars indicate expression levels in aged sham, white bars adult sham. Light blue bars represent iTBS and dark blue bars acc-iTBS. **(A)** iTBS and acc-iTBS show no effect in the aged cerebellum. **(B)** BHFS has small effect on adult has represented by pink bars. *p < 0.05, ****p < 0.0001.

**Supplementary Fig. 2. iTBS decreases developmental biological processes in adult cerebellum.**

**(A)** Gene Ontology (GO) of biological processes decreased by iTBS in adult. The left vertical axis lists the top biological processes identified through GO Analysis (using GSEA website). Each pathway is associated with a gene count on the x-axis (number of altered genes in that pathway) and an adjusted p value gradient, indicating the significance of enrichment. Bubble size reflects the proportion of downregulated genes included in each biological process. **(B)** GO map of biological processes downregulated by iTBS in adult. CLueGO results display a functionally grouped network of pathways. The most significant pathway in each cluster is designated as the leading term. Node size represents pathway significance, and interactions between pathways are indicated.

**Supplementary Fig. 3. Purkinje Cell morphology is not affected by acute LI-rTMS.**

**(Ai)** Dendritic height and density **(Aiii)** were the same at both ages (Height: Two-way ANOVA F_1,35=_0.0419 p=0.839; stimulation F_1,35 =_ 5.925 p=0.02; Density: Two-way ANOVA F_1,35=_0.5631 p=0.458; stimulation F_1,35 =_ 4.404 p=0.043). Acute LI-rTMS globally decreased height and density even if no specific effects were observed in either adults or aged (Bonferroni post –hoc p>0.05). **(Aii)** width was not affected by age or stimulation (Two-way ANOVA F_1,35=_0.0016 p=0.9686; stimulation F_1,35 =_ 0.1594 p=0.6921). **(B)** Sholl analysis, quantifying the number of dendritic branches as function of distance from the soma, does not show any significant effect of short -term LI-rTMS on Purkinje dendritic tree complexity (Two-way ANOVA and pairwise comparisons, Bonferroni correction, all p>0.05). Intergroup comparisons: *p≤0.05

**Supplementary Fig. 4. All animals can orientate themselves and swim to a platform.**

There were no significant differences between any groups for swim speed **(Ai),** distance travelled **(Aii)** or the latency to the platform **(Aiii)** (Kruskal-Wallis test, all p>0.05).

**Supplementary Fig. 5. RNA SEQ analysis workflow.**

The right and left cerebella were harvested 24 hours after the last stimulation and the right cerebella were used for the following steps. Total RNA was extracted using the TRIzol reagent (Ambion) according to the manufacturer’s instructions. The overall RNA quality was assessed using Agilent High Sensitivity RNA ScreenTape System. Sequencing was performed using Illumina NovaSeq (paired-end sequencing, 800 million reads) by Next Generation Sequencing Platform (NGS) (Institut du Cerveau, Paris). The reference mouse genome (mm10) was downloaded in compressed FASTA format from UCSC’s genomes (University of California, Santa Cruz). Furthermore, the comprehensive gene annotation for mm10 was downloaded in the GTF format from GENCODE (Release M14; GRCm38.p5). Paired-end read files were aligned to version mm10 of mouse reference genome and transcriptome using HiSat2 (v2.2.1). Quality control checks were assessed using FastQC (v0.74) and summarized in a single report generated by MultiQC (v1.11). The number of aligned reads was counted by featurecounts tool (v2.0.3). Finally, we used the DESeq2 (v2.11.40.8) package to determine differentially expressed genes (DEG) from count tables. In the present study, genes with adjusted p value pAdj<0.05 were selected as significant. Volcano plots were generated to show the significantly upregulated and downregulated genes. Further analysis and data visualisation were performed using the Python v3.12.13 package. Functional enrichment analysis was performed using the Cytoscape (v3.10.3) software and the ClueGo (v2.5.10) extension to identify Gene Ontology (GO) term enrichment (Biological Processes) amongst DEGs with a p-value adjustment (pAdj) threshold of 0.05. ClueGo also allowed visualization of clusters of pathways/terms. The parameters are specified in the supplementary Table 1.

**Supplementary Table 1. ClueGo parameters on Cytoscape**

**Supplementary Table 2. qPCR Primers**

