## Supplementary files for "Age-related cerebellar genetic, neuronal and functional impairments are reversed by specific magnetic stimulation protocols"

### Supplementary files for Fauquier et al

**Supplementary Figure 1. LI-rTMS has different effects depending on age and pattern.**

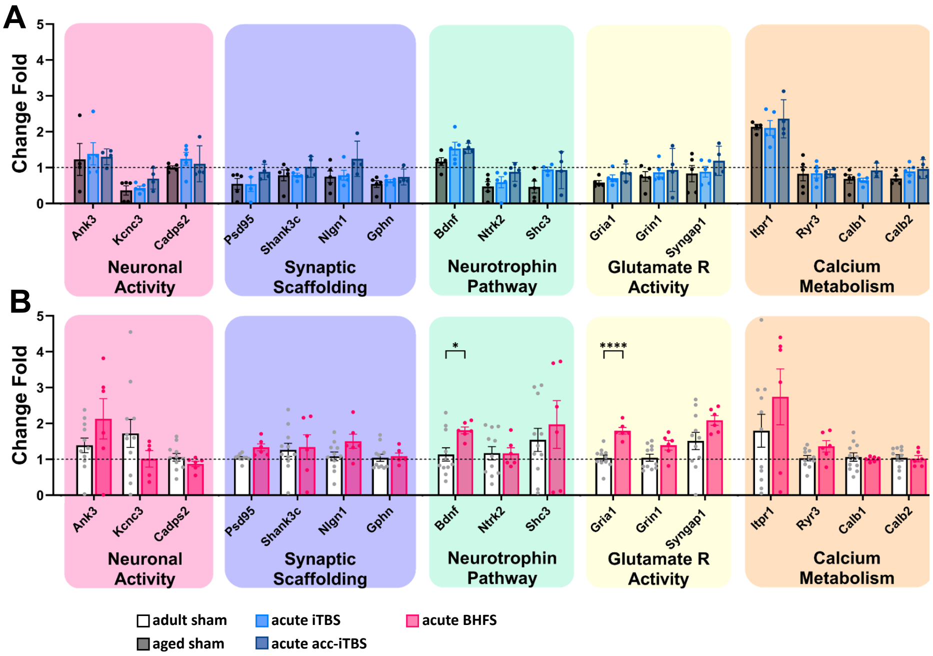


Gene expression data are expressed as fold change or relative expression normalized to the adult sham group, according to the Pfaffl equation. The horizontal dotted line represents the mean expression (geometric mean) of adult sham animals. Dark grey bars indicate expression levels in aged sham, white bars show adult sham. Light blue bars represent iTBS and dark blue bars acc-iTBS. **(A)** iTBS and acc-iTBS show no effect in the aged cerebellum. **(B)** BHFS has small effect in the adult cerebellum.

*p < 0.05, ****p < 0.0001.

**Supplementary Figure 2. iTBS decreases developmental biological processes in adult cerebellum.**

A

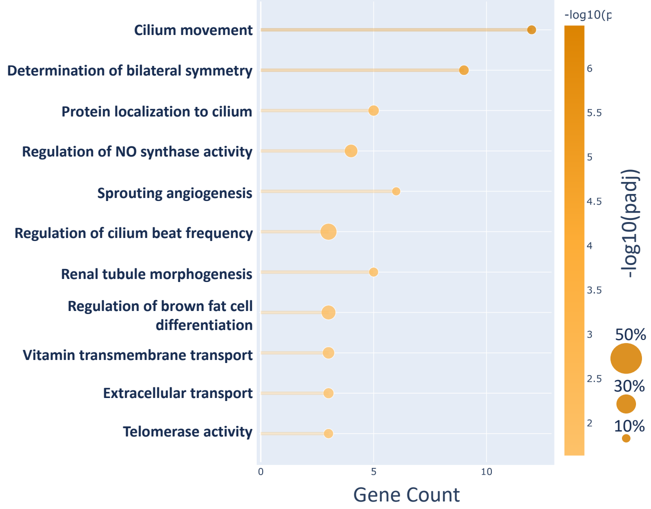


B

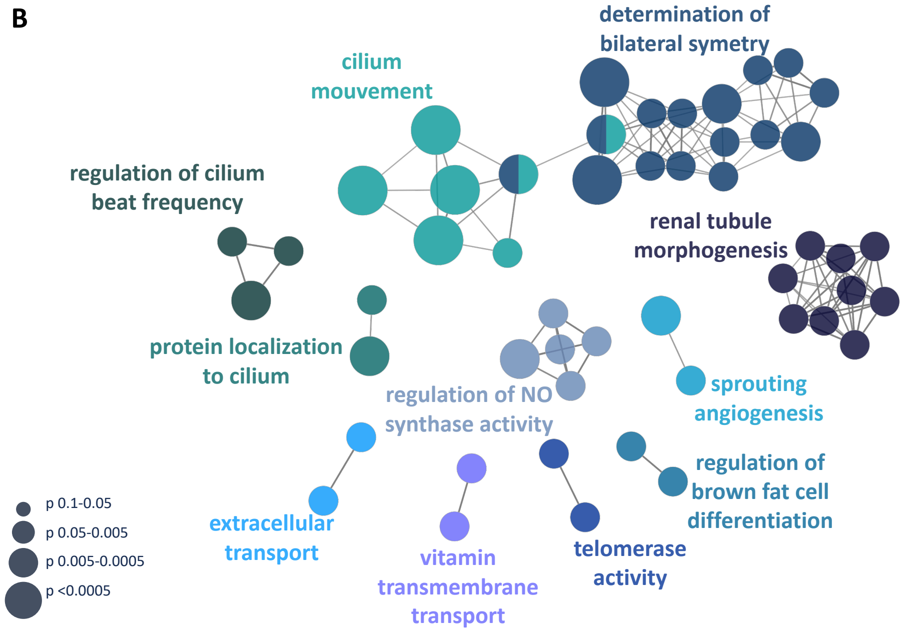


**(A)** Gene Ontology (GO) of biological processes decreased by iTBS in adult. The left vertical axis lists the top biological processes identified through GO Analysis (using GSEA website). Each pathway is associated with a gene count on the x-axis (number of altered genes in that pathway) and an adjusted p value gradient, indicating the significance of enrichment. Bubble size reflects the proportion of downregulated genes included in each biological process. **(B)** GO map of biological processes downregulated by iTBS in adult. CLueGO results display a functionally grouped network of pathways. The most significant pathway in each cluster is designated as the leading term. Node size represents pathway significance, and edges indicate interactions between pathways.

**Supplementary Figure 3. Purkinje Cell morphology is not specifically affected by acute LI-rTMS.**

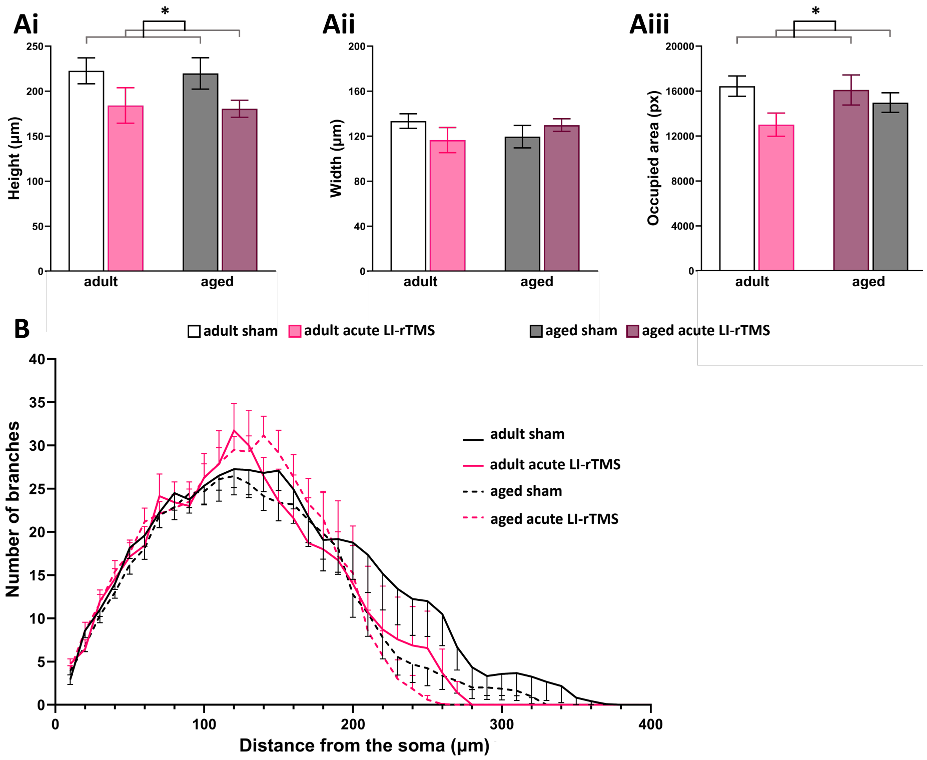


**(Ai)** Dendritic height and density **(Aiii)** were the same at both ages (Height: Two-way ANOVA $F_{1,35}$=0.0419 p=0.839; stimulation $F_{1,35}$ = 5.925 p=0.02; Density: Two-way ANOVA $F_{1,35}$=0.5631 p=0.458; stimulation $F_{1,35}$ = 4.404 p=0.043). Acute LI-rTMS globally decreased height and density even if no specific effects were observed in either adults or aged (Bonferroni post –hoc p>0.05). **(Aii)** width was not affected by age or stimulation (Two-way ANOVA $F_{1,35}$=0.0016 p=0.9686; stimulation $F_{1,35}$ = 0.1594 p=0.6921). **(B)** Sholl analysis, quantifying the number of dendritic branches as function of distance from the soma, does not show any significant effect of short -term LI-rTMS on Purkinje dendritic tree complexity (Two-way ANOVA and pairwise comparisons, Bonferroni correction, all p>0.05).

Intergroup comparisons: *p≤0.05

**Supplementary Figure 4. All animals can orientate themselves and swim to a visible platform.**

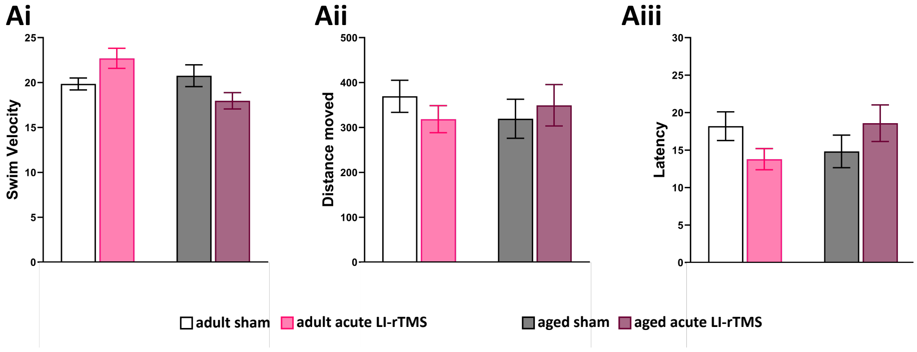


In the visible platform test of the Morris water maze, there were no significant differences between any groups for swim speed **(Ai),** distance travelled **(Aii)** or the latency to the platform **(Aiii)** (Kruskal-Wallis test, all p>0.05), showing that all animals could orientate and swim to a specific location.

**Supplementary Figure 5. RNA SEQ analysis workflow.**

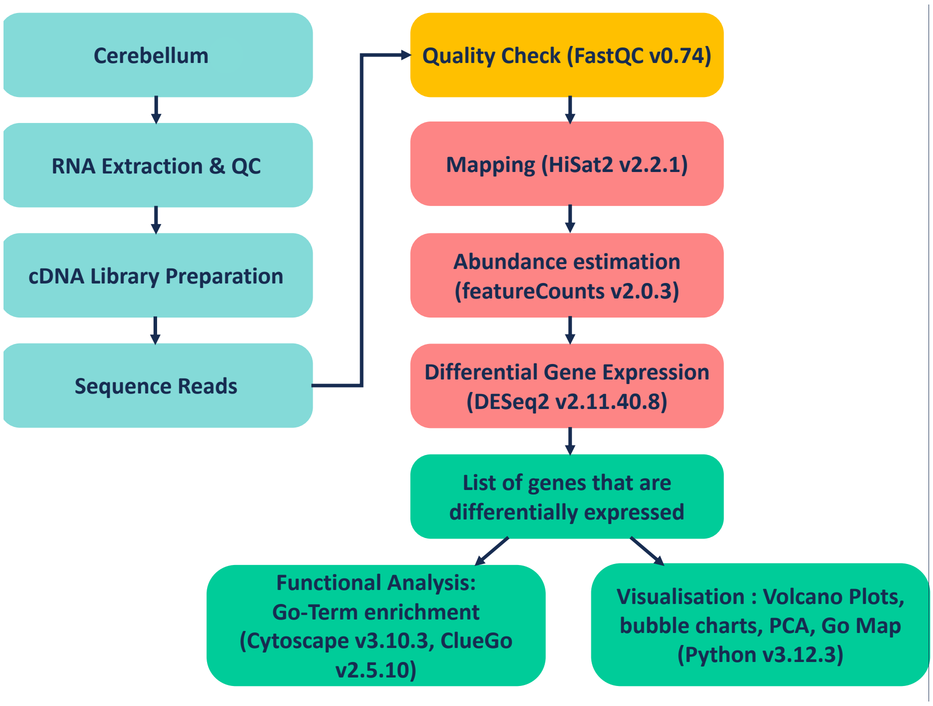


The right and left hemicerebella were harvested 24 hours after the last stimulation and the right hemicerebella were used for the following steps. Total RNA was extracted using the TRIzol reagent (Ambion) according to the manufacturer's instructions. The overall RNA quality was assessed using Agilent High Sensitivity RNA ScreenTape System. Sequencing was performed using Illumina NovaSeq (paired-end sequencing, 800 million reads) by Next Generation Sequencing Platform (NGS) (Institut du Cerveau, Paris). The reference genome for mouse (mm10) was downloaded in compressed FASTA format from UCSC's genomes (University of California, Santa Cruz). Furthermore, the comprehensive gene annotation for mm10 was downloaded in the GTF format from GENCODE (Release M14; GRCm38.p5). Paired-end read files were aligned to version mm10 of mouse reference genome and transcriptome using HiSat2 (v2.2.1). Quality control checks were assessed using FastQC (v0.74) and summarized in a single report generated by MultiQC (v1.11). The number of aligned reads was counted by featurecounts tool (v2.0.3). Finally, we used the DESeq2 (v2.11.40.8) package to determine differentially expressed genes (DEG) from count tables. In the present study, genes with adjusted p value pAdj<0.05 were selected as significant. Volcano plots were generated to show the significantly upregulated and downregulated genes. Further analysis and data visualisation were performed using the Python v3.12.13 package. Functional enrichment analysis was performed using the Cytoscape (v3.10.3) software and the ClueGo (v2.5.10) extension to identify Gene Ontology (GO) term enrichment (Biological Processes) amongst DEGs with a p-value adjustment (pAdj) threshold of 0.05. ClueGo also allowed visualization of clusters of pathways/terms. The parameters are specified in the supplementary Table 1.

#### Supplementary Table 1. ClueGo parameters on Cytoscape

| | Sham18-4 Up DEG | Itbs4 vs Sham4 Up DEG | Itbs4 vs Sham4 Down DEG | Bhfs18 vs Sham18 Down DEG | Bhfs18 vs Sham18 Up DEG |
|---|---|---|---|---|---|
| Prefiltering on DEG | pAdj <0,05<br>foldchange > 0 | pAdj <0,05<br>foldchange > 0 | pAdj <0,05<br>foldchange > 0 | pAdj <0,05<br>foldchange < -0.4 | pAdj <0,05<br>foldchange > 0.4 |
| Ontologies/ Patyways | Go-Biological Process (25/05/22) | Go-Biological Process (25/05/22) | Go-Biological Process (25/05/22) | Go-Biological Process (25/05/22) | Go-Biological Process (25/05/22) |
| Network Specificity | Medium | Medium | Medium | Medium | Medium |
| Use GO Term Fusion | Yes | No | No | Yes | Yes |
| Show only Pathways with pV <=0,05 | Yes | Yes | Yes | Yes | Yes |
| **Advanced Term Selection Options** | | | | | |
| Go Tree interval | 1 to 3 | All | All | 0 to 4 | 0 to 4 |
| Cluster | 3 Min #Genes and 4%Genes | 3 Min #Genes and 4%Genes | 3 Min #Genes and 4%Genes | 3 Min #Genes and 4%Genes | 3 Min #Genes and 4%Genes |
| Go Term Network Connectivity (Kappa Score) | 0,4 | 0,4 | 0,4 | 0,4 | 0,4 |
| Statistical Options | Default (Bonferroni step down) | Default (Bonferroni step down) | Default (Bonferroni step down) | Default (Bonferroni step down) | Default (Bonferroni step down) |
| **Grouping Options** | | | | | |
| Use Go Term Grouping | Yes | Yes | Yes | Yes | Yes |
| Leading Group Term based on | Highest Significance | Highest Significance | Highest Significance | Highest Significance | Highest Significance |
| Initial Group Size | 2 | 1 | 2 | 2 | 2 |
| %Genes for Group Merge | 35 | 50 | 50 | 30 | 30 |
| %Term for Group Merge | 35 | 50 | 50 | 30 | 30 |

#### Supplementary Table 2. qPCR Primers

**Genes of Interest**

| | Forward | Tm | Reverse | Tm |
|---|---|---|---|---|
| *Ank3* | TGGAAACCACACAGCGGAAGTC | 62,1 | TTTCATGCCGACAGGCACACAG | 62,1 |
| *Bdnf* | ACTATGGTTATTTCATACTTCGGTTGC | 60,0 | TCAGCCAGTGATGTCGTCG | 59,8 |
| *Cadps2* | CCAGAATGGTTCAAAGTGGAGG | 59,2 | GGCTACGGACACGTTTTTCTATG | 59,9 |
| *Calb1* | CATCTCTGATCACAGCCTCACA | 59,8 | GATAGCTCCAATCCAGCCTTCT | 59,6 |
| *Calb2* | GAAAATTGAGATGGCGGAGCTG | 60,2 | GTCATACTTCCGCCAAGCCTT | 60,7 |
| *Gphn* | ATCGCATGTCTCCTTTTCCCC | 60,4 | TGTAACCCGCATCACTTGTCC | 60,6 |
| *Gria1* | CTAGGCTGCCTGAACCTTTGG | 61,0 | GGGGAAGATTGAATGGAAGCAG | 59,3 |
| *Grin1* | ATCATCTGGCCAGGAGGAGAGA | 62,1 | CCATCACTCATTGTGGGCTTGA | 60,9 |
| *Itrpr1* | GACATCCTGATTGAGACCAAGC | 59,1 | CTGTCCCTCTTTAGCATCTTGC | 59,1 |
| *Kcnc3* | CTCCTCTAGCTAGGCTGGGTC | 60,8 | GAGGCAGACGTGATTGGACAG | 61,0 |
| *Nlgn1* | GGGATGAGGTTCCCTATGTGTT | 59,5 | GGTTGGGTTTGGTATGGATGAA | 58,6 |
| *Ntrk2* | CCATTTAACTGCACCCGCAC | 59,4 | CCCAAGACCAGCAGGCATAA | 59,4 |
| *Psd95* | ACTGCATCCTTGATGTCTCAGC | 60,2 | TTCGATGACACGTTTCACTTTGT | 57,1 |
| *Ryr3* | AATCCACAGCCTTCTCCTTCC | 59,7 | GAACAAAGCAGACAGAATCCCC | 59,5 |
| *Shank3c* | TCTGTGACCAGGAAACCCGA | 60,8 | CTCCCCTTTTCTTCTCCGAATGG | 60,9 |
| *Shc3* | AGGGAAGAGCAAGCCATTGCAC | 62,1 | AAAACGACATCCACCCGCTGAC | 62,1 |
| *Syngap1* | GCTACACATGTCCAACCGGA | 60,0 | AGCCTGCCAATGATGCTCTT | 60,0 |

**Housekeeper genes**

| | Forward | Tm | Reverse | Tm |
|---|---|---|---|---|
| *Hprt* | CGTTTCTGAGCCATTGCTGAG | 59,8 | TCATCGCTAATCACGACGCTG | 59,8 |
| *Mrpl32* | CGTTGCTGCTGCTTTCACTAC | 59,8 | ACTGACAAAGCGACTCCAGCT | 59,8 |
